# Glutamine metabolism enables NKT cell homeostasis and function through the AMPK-mTORC1 signaling axis

**DOI:** 10.1101/2021.10.07.463490

**Authors:** Ajay Kumar, Emily L. Yarosz, Anthony Andren, Li Zhang, Costas A. Lyssiotis, Cheong-Hee Chang

**Affiliations:** Department of Microbiology and Immunology, University of Michigan Medical School, Ann Arbor, MI 48109, USA; Immunology Graduate Program, University of Michigan Medical School, Ann Arbor, MI 48109, USA; Department of Molecular and Integrative Physiology, University of Michigan Medical School, Ann Arbor, MI 48109, USA; Department of Internal Medicine, Division of Gastroenterology and Hepatology, University of Michigan, Ann Arbor, MI 48109, USA; Rogel Cancer Center, University of Michigan, Ann Arbor, MI 48109, USA

**Keywords:** Metabolism, glutathione, ROS, glycosylation

## Abstract

Cellular metabolism is essential in dictating conventional T cell development and function, but its role in natural killer T (NKT) cells has not been well studied. We have previously shown that NKT cells operate distinctly different metabolic programming from CD4 T cells, including a strict requirement for glutamine metabolism to regulate NKT cell homeostasis. However, the mechanisms by which NKT cells regulate glutamine metabolism for their homeostasis and effector functions remain unknown. In this study, we report that steady state NKT cells have higher glutamine levels than CD4 T cells and NKT cells increase glutaminolysis upon activation. Among its many metabolic fates, NKT cells use glutamine to fuel the tricarboxylic acid cycle and glutathione synthesis, and glutamine-derived nitrogen enables protein glycosylation via the hexosamine biosynthesis pathway (HBP). Each of these functions of glutamine metabolism was found to be critical for NKT cell survival and proliferation. Furthermore, we demonstrate that glutaminolysis and the HBP differentially regulate IL-4 and IFNγ production. Finally, glutamine metabolism appears to be controlled by AMP-activated protein kinase (AMPK)-mTORC1 signaling. These findings highlight a unique metabolic requirement of NKT cells which can be potentially serve as an effective immunotherapeutic agent against certain nutrient restricted tumors.

**Significance:** NKT cells get activated very early during an immune response and produce cytokines and chemokines, which further activate other immune cell types. Although metabolism regulates these functions in other T cell subsets, little is understood about how metabolic pathways are controlled in NKT cells. The present study shows that NKT cells metabolize the amino acid glutamine through two different branches of metabolism, which control NKT cell homeostasis and expansion in a similar manner but control cytokine production differently. This glutamine dependency seems to be regulated by AMP-activated protein kinase (AMPK), which is a central regulator of energy homeostasis. Together, our study demonstrates a unique metabolic profile of glutamine metabolism in NKT cells which could be harnessed for NKT cell-based immunotherapy.

## Introduction

Cellular metabolism plays a significant role in modulating T cell functions. Activated T cells undergo metabolic rewiring to fulfill the demands of clonal expansion as well as cytokine synthesis and secretion. A recent body of evidence has highlighted the role of cellular metabolism in regulating T cell plasticity. T cells shift glucose metabolism from a more glycolytic phenotype to a more oxidative phenotype after activation, a process known as metabolic reprogramming (1-3). This metabolic reprogramming is orchestrated by a series of signaling pathways and transcriptional networks (4-6). Additionally, the various T cell subsets operate distinct metabolic profiles that are critical for their specific effector functions (6, 7).

Invariant natural killer T (NKT) cells are innate-like lymphocytes that recognize glycolipid antigens in the context of the nonclassical MHC molecule CD1d, which is present on antigen presenting cells. NKT cells are selected by cortical thymocytes expressing CD1d and mature through a series of stages (8, 9). Thymic NKT cells are capable of producing the cytokines IFNγ, IL-4, and IL-17 and are thus termed NKT1, NKT2, and NKT17, respectively (10). NKT cells are a vital part of the defense against infectious diseases (11-13) and also play a role in the development of autoimmunity (14, 15) and asthma (16). Additionally, NKT cells mediate potent antitumor immune responses and have been utilized in immunotherapy for cancer patients using various immunomodulatory approaches (17-20).

NKT cells express promyelocytic leukemia zinc finger (PLZF, encoded by *Zbtb1*6), a transcription factor required for NKT cell development and function (8, 21, 22). Several studies have shown that metabolic signals are critical for NKT cell development and function. Mammalian target of rapamycin (mTOR) complex 1 and complex 2 integrate various environmental cues to regulate cellular growth, proliferation, and metabolism (23, 24). Deletion of either mTORC1 or mTORC2 leads to a block in NKT cell development during which NKT cells accumulate in the early developmental stages (25, 26). Additionally, mTORC1 is a critical regulator of glycolysis and amino acid transport in T cells (27, 28). mTORC1 has been shown to be negatively regulated by AMP-activated protein kinase (AMPK) in T cells (29). AMPK senses cellular energy levels and in turn activates pathways necessary to maintain cellular energy balance. Additionally, loss of the AMPK-interacting adaptor protein folliculin-interacting protein 1 (Fnip1) results in defective NKT cell development (30).

As NKT cells develop and mature in the thymus, they become more quiescent and display lower metabolic activity in the peripheral organs compared to conventional T cells (31). We have shown that resting NKT cells have lower glucose uptake and mitochondrial function compared to conventional T cells, which is regulated by PLZF (32). Furthermore, high environmental levels of lactate are detrimental for NKT cell homeostasis and cytokine production, suggesting that reduced glycolysis is essential for NKT cell maintenance (32). Interestingly, NKT cells preferentially partition glucose into the pentose phosphate pathway (PPP) and contribute less carbon into the tricarboxylic acid (TCA) cycle than CD4 T cells. Recently, lipid synthesis has also emerged as a critical regulator of NKT cell responses (33).

In addition to glucose, rapidly proliferating cells require the amino acid glutamine to produce ATP, biosynthetic precursors, and reducing agents (6, 34). Glutaminolysis refers to the breakdown of the glutamine to fuel metabolism. In some proliferating cell types, glutaminolysis can take place in the mitochondria, where glutamine is converted to glutamate by the glutaminase (GLS) enzyme. From here, glutamate can undergo several metabolic fates. For one, glutamate can be deaminated into the TCA cycle intermediate α-ketoglutarate (αKG) by either glutamate dehydrogenase (GDH) or aminotransferases. Glutamate can also be transported back into the cytosol and produce glutathione (GSH), a critical mediator of cellular redox balance (35). Additionally, glutamine-derived nitrogen can be used to fuel *de novo* glycosylation precursor biogenesis in the hexosamine biosynthesis pathway (HBP) (36, 37).

A growing body of work has recently begun to highlight the importance of glutamine metabolism in modulating T cell-mediated immunity. Activated T cells not only upregulate amino acid transport but also increase the expression of enzymes involved in glutamine metabolism (6, 34). In addition, glutamine deprivation suppresses tumor growth and induces cell death in several cancer types (38, 39). The glutamine dependency displayed by cancerous cells has been referred to as glutamine addiction (40, 41). Similarly, we have previously shown that NKT cells rely on glutamine for their survival and proliferation (32). Despite this, the precise metabolic pathways and outputs of glutamine metabolism in NKT cells remain unknown.

In the current study, we report that NKT cells have higher glutamine metabolism than CD4 T cells, and NKT cells enhance glutamine metabolism after activation. NKT cells use glutamine-derived carbon to fuel the TCA cycle and glutamine-derived nitrogen to fuel the HBP while simultaneously supporting GSH generation via glutamine-derived glutamate. More importantly, these processes are critical for NKT cell survival and proliferation. NKT cells require glutaminolysis for IL-4 production, but they use the HBP to support IFNγ production. Furthermore, we demonstrate that NKT cells are glutamine addicted, as glucose is not sufficient to fuel mitochondrial function in the absence of glutamate oxidation. Lastly, AMPK-mTORC1 signaling is involved in the regulation of glutamine metabolism in NKT cells.

## Results

### NKT cells upregulate glutamine metabolism upon activation

We previously reported that resting NKT cells are less glycolytic than CD4 T cells and rely on glutamine for their survival and proliferation (32). To gain a better understanding of glutamine metabolism in NKT cells, we assessed metabolite levels in freshly sorted NKT and CD4 T cells using liquid chromatography (LC)-coupled tandem mass spectrometry (LC-MS/MS)-based metabolomics. Metabolomic analysis showed that NKT cells have lower levels of metabolites related to glycolysis but higher levels of metabolites related to glutaminolysis compared to CD4 T cells (Fig. 1A). Pathway enrichment analysis revealed increased amino acid metabolism in NKT cells, which includes glutamine metabolism (Fig. S1A). In addition to glutamine, other metabolites such as glutamate, arginine, and asparagine were relatively high in NKT cells compared to CD4 T cells (Fig. 1B). To investigate whether NKT cells upregulate glutaminolysis upon activation, we measured intracellular metabolite levels after 3 days of stimulation using LC-MS/MS. Metabolites from the culture media were measured simultaneously. Metabolites downstream of glutamine metabolism were increased and decreased in cells and culture media, respectively, upon activation (Fig. 1C and 1D). These data suggest that NKT cells enhance both glutamine import and utilization during activation. Indeed, the expression of CD98, a heterodimeric amino acid transporter known to uptake glutamine (42), was increased on activated NKT cells (Fig. S1B). Moreover, the levels of metabolites derived from glutamine such as glutamate, αKG, and GSH were increased after activation (Fig. 1C, and S1C-S1E). We also observed that the expression of genes encoding key enzymes involved in glutamine metabolism was elevated after activation (Fig. S1F). Overall, NKT cells upregulate glutamine metabolism upon activation.

**Fig. 1.**
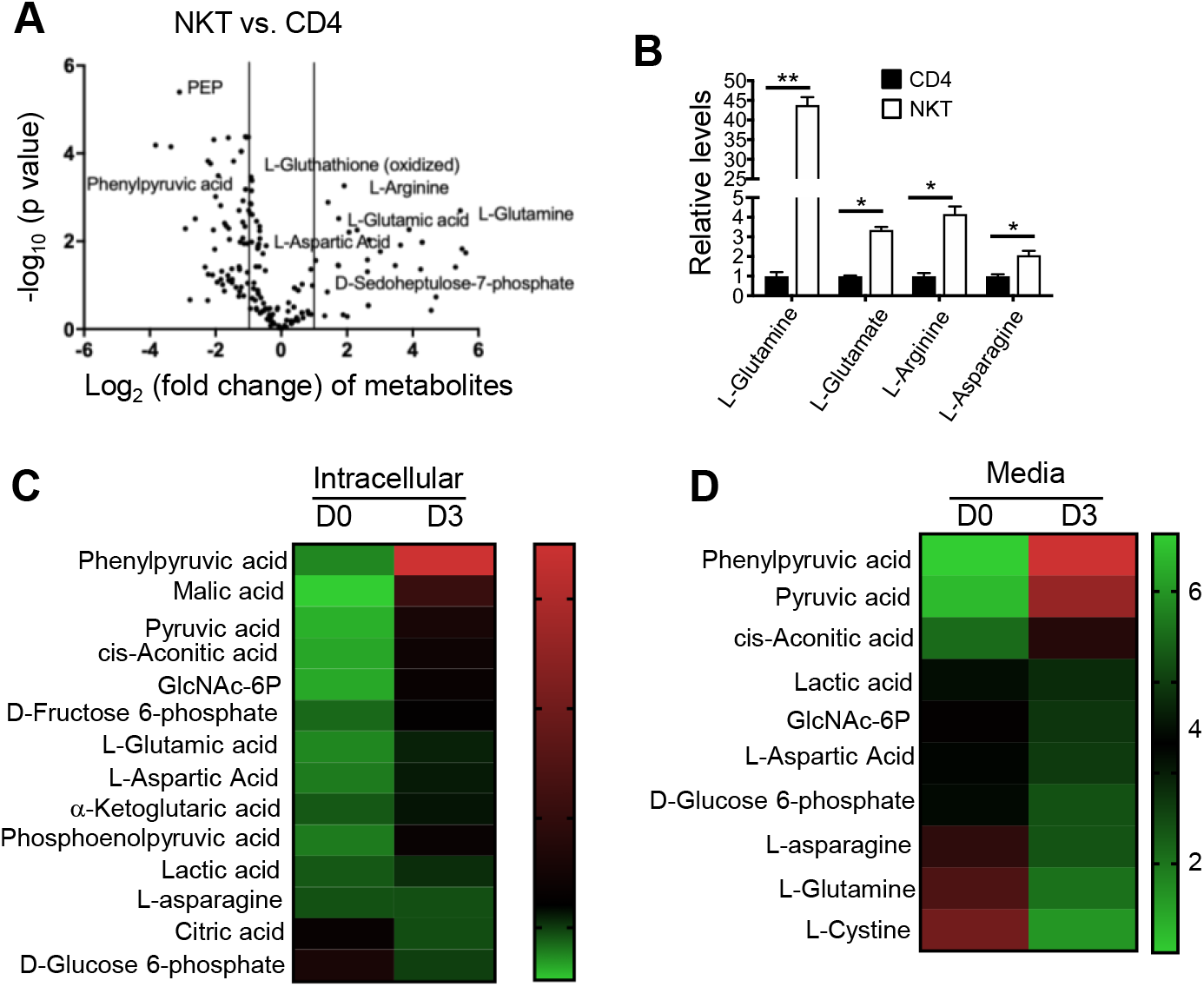
NKT cells increase glutaminolysis upon activation. (A and B) Freshly sorted NKT and CD4 T cells from C57BL/6 mice were subjected to metabolomic analysis through LC-MS/MS. (A) The volcano graph depicts upregulated and downregulated metabolites in resting NKT cells compared to CD4 T cells (n=3). (B) Graph shows relative levels of the indicated metabolites in resting NKT cells vs. CD4 T cells (n=3). (C and D) NKT cells were stimulated with αGalCer (100 ng/ml) for 3 days. The cell lysate was prepared from unstimulated (D0) and stimulated (D3) NKT cells. Media was also collected on day 3 (D3) of activation. Cell lysate and media samples were subjected to metabolomic analysis through LC-MS/MS. (C) Heatmap represents relative levels of metabolites in unstimulated and stimulated NKT cells (n=3). (D) Heatmap shows relative levels of metabolites in the media collected from unstimulated and stimulated NKT cells (n=3). Data are shown as mean ± SEM. *p<0.05, **p<0.01 were considered significant.

### Glutaminolysis is essential for NKT cell survival and proliferation

Glutamine is a major source of energy and carbon molecules in rapidly proliferating cells like immune cells and cancerous cells (41). NKT cells have been shown to rely on glutamine for their survival and proliferation (32), prompting us to investigate whether this dependency on glutamine is due to glutaminolysis. We used a variety of pharmacological inhibitors to examine the importance of each branch of glutamine catabolism for NKT cell responses (Fig. 2A). To begin, we measured glutamate in NKT cells activated under glutamine deprivation conditions. We found that glutamate levels are decreased when cells are grown in the absence of glutamine (Fig. 2B). Next, to confirm whether the oxidation of glutamine into glutamate is necessary for NKT cell survival and proliferation, cells were activated in the presence or absence of the GLS inhibitor CB839. GLS inhibition impaired NKT cell survival, proliferation, and activation (Fig. 2C).

**Fig. 2.**
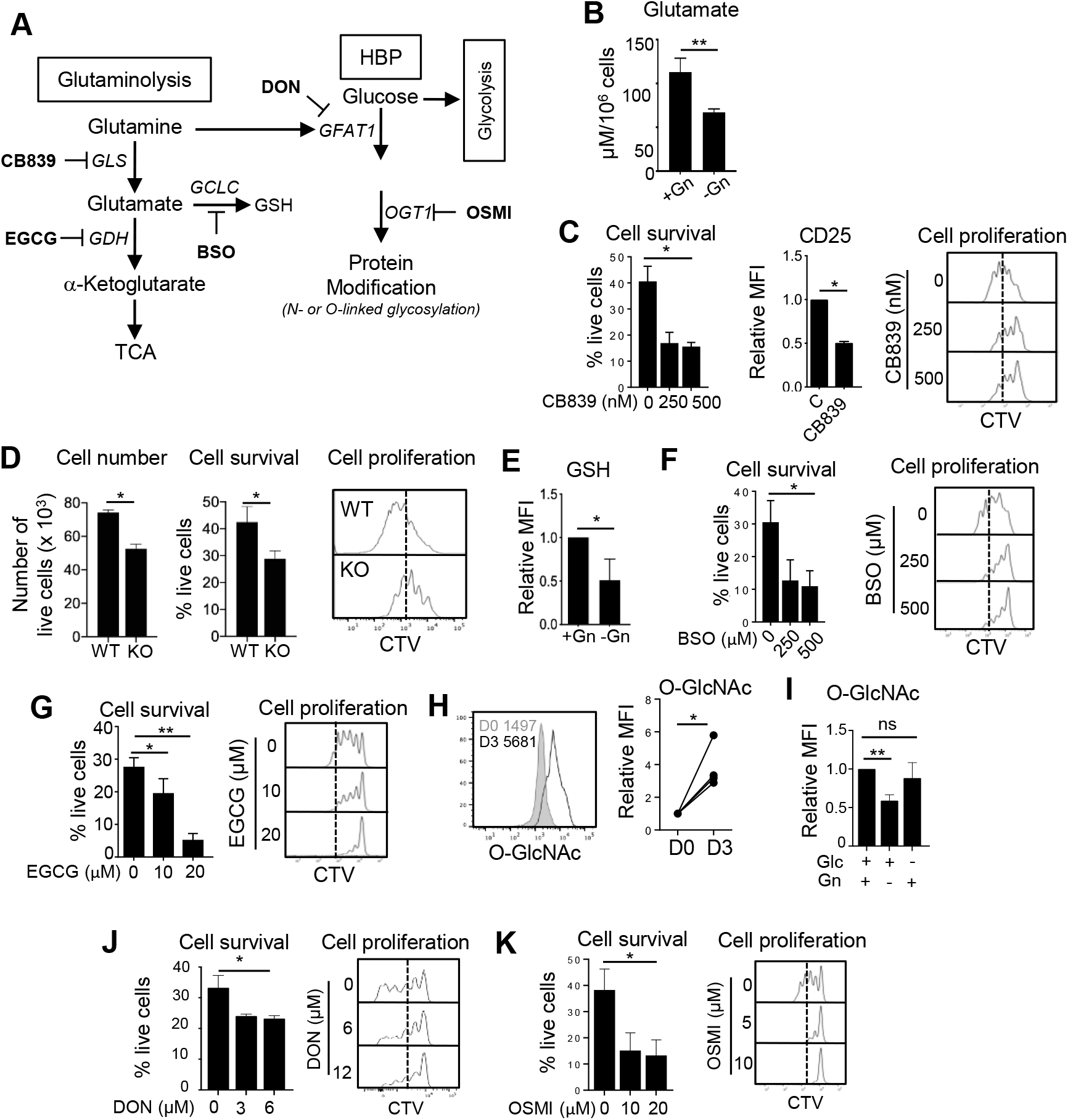
Glutamine metabolism is essential for NKT cell survival and proliferation. (A) The schematic depicts key branches of glutamine metabolism producing αKG and GSH as well as utilization of glutamine in the HBP to synthesize O-GlcNAc. Pathway-specific inhibitors are shown in bold and the names of target enzymes are italicized. (B and C) Sorted NKT cells from C57BL/6 mice were labeled with 5 μM CellTrace Violet (CTV) and stimulated with αGalCer (100 ng/ml) for 3 days in the indicated culture conditions. (B) The graph shows glutamate levels in NKT cells activated in the presence or absence of glutamine (n=3). (C) Graphs show cell survival (relative levels of percentages of live cells) measured by live/dead marker staining, CD25 expression, and cell proliferation of NKT cells activated in the presence or absence of CB839 (n=4). Control levels were set at 1. (D) NKT cells from WT and GLS1 KO mice were activated for 3 days as in (B). Graphs show total live NKT cell numbers, percentages of cell survival, and cell proliferation (n=3). (E) Sorted NKT cells were activated in the presence or absence of glutamine. GSH levels on day 3 of activation are shown (n=3). (F and G) NKT cells were activated for 3 days in the presence or absence of BSO (F) or EGCG (G). Graphs show cell survival and proliferation (n=3). (H) The levels of O-GlcNAc in NKT cells with and without activation were compared (n=3). (I) Sorted NKT cells were stimulated for 3 days in the presence or absence of either glutamine or glucose as indicated. The graph shows the relative mean fluorescent intensity (MFI) of O-GlcNAc on day 3 of activation (n=3). (J and K) NKT cells were activated for 3 days in the presence or absence of DON (J) or OSMI (K). Cell survival and proliferation were shown (n=3). All data are representative of or combined from at least three independent experiments. Data are shown as mean ± SEM. *p<0.05, **p<0.01.

Next, we used mice having a T cell-specific deletion of GLS1 (GLS1^fl/fl^ CD4-Cre, referred to as GLS1 KO) (6) to validate the responses caused by the pharmacological inhibitor. GLS1 deficiency did not affect NKT cell development in the thymus, but NKT cell numbers were slightly reduced in the spleens of these mice (Fig. S2A and S2B), suggesting a role of glutamine in peripheral NKT cell maintenance. Next, we measured cell survival and proliferation in activated WT and GLS1 KO NKT cells. Similar to what was seen with CB839, GLS1 deficient cells not only died more than WT cells but also proliferated worse than WT cells (Fig. 2D).

Because glutamine contributes to cellular redox regulation through glutathione (GSH) synthesis, we investigated whether glutamine is converted to GSH in the absence of glutamine. As expected, GSH levels were decreased in NKT cells grown under glutamine deprivation conditions (Fig. 2E). GSH is critical for NKT cell homeostasis, since cell survival and proliferation were impaired when GSH synthesis was inhibited by adding buthionine sulfoximine (BSO) to the culture media (Fig. 2F).

In addition to GSH, αKG is produced from glutamate, after which it can then enter the TCA cycle. The elevated levels of αKG in activated NKT cells prompted us to examine if the conversion of glutamate to αKG is critical for NKT cells. Like GLS inhibition, GDH inhibition using the pan dehydrogenase inhibitor epigallocatechin-3-gallate (EGCG) reduced cell survival and proliferation (Fig. 2G). Furthermore, GDH inhibition decreased mitochondrial mass and mitochondrial membrane potential (Fig. S2C). To confirm whether glutamate contributes to mitochondrial energy production in NKT cells, ATP was measured. ATP levels decreased significantly after GLS inhibition, suggesting that glutamate fuels mitochondrial ATP production (Fig. S2D). Therefore, glutaminolysis is necessary not only for mitochondrial anaplerosis but also for GSH synthesis and ATP production. Collectively, glutaminolysis is critical for NKT cell survival and optimal proliferation, potentially by supporting mitochondrial function.

### NKT cell homeostasis depends upon the contribution of glutamine-derived nitrogen to the hexosamine biosynthesis pathway

Glucose and glutamine contribute carbon and nitrogen, respectively, via the HBP in T cells to generate UDP-GlcNAc, the primary donor for cellular glycosylation (Fig. 2A) (37). The HBP deposits O-linked and N-linked glycosylation marks on proteins, which are necessary for protein stability and function. To test the role of *de novo* glycosylation biosynthesis via the HBP in NKT cells, we examined total protein glycosylation upon glutamine deprivation by measuring O-GlcNAc-ylation of the proteome. Activated NKT cells showed increased total protein glycosylation (Fig. 2H) as well as higher mRNA expressions of both the *Gfat*1 and *Ogt* genes (Fig. S2E). Both glucose and glutamine are required for HBP initiation. Next, to understand how nutrient limitation impacts the HBP, O-GlcNAc levels were measured in cells stimulated in the presence of glucose only, glutamine only, or both. We found that glutamine limitation reduced *de novo* O-GlcNAc synthesis significantly more than glucose limitation does in activated NKT cells (Fig. 2I) suggesting increased salvage pathway for HBP synthesis under glucose restriction.

To determine the role of the HBP in NKT cell responses, we treated NKT cells with 6-diazo-5-oxo-L-nor-leucine (DON), a pan glutamine-deamidase inhibitor (43), during activation. We observed that DON treatment reduced O-GlcNAc levels (Fig. S2F) leading to impaired NKT cell survival accompanied by reduced cell proliferation (Fig. 2J). Similarly, inhibition of OGT by OSMI resulted in more cell death and less cell proliferation than untreated cells (Fig. 2K). These data suggest that the HBP is essential for NKT cell homeostasis.

### NKT cell homeostasis requires GSH-mediated redox balance

NKT cells rely on glutamine for GSH, which is vital for the effective management of reactive oxygen species (ROS) (44). Therefore, NKT cells may be susceptible to cell death in the absence of GSH. We have previously shown that NKT cells are highly susceptible to oxidative stress (45). Since GSH maintains intracellular redox balance, we examined total ROS production in NKT cells treated with the GSH inhibitor BSO. Total ROS levels, as measured by DCFDA, were greater in the presence of the inhibitor than the control (Fig. 3A). In contrast, GSH inhibition reduced mitochondrial ROS, mitochondrial mass, and mitochondrial potential (Fig. 3B-3D), suggesting that GSH is critical for mitochondrial functions.

**Fig. 3.**
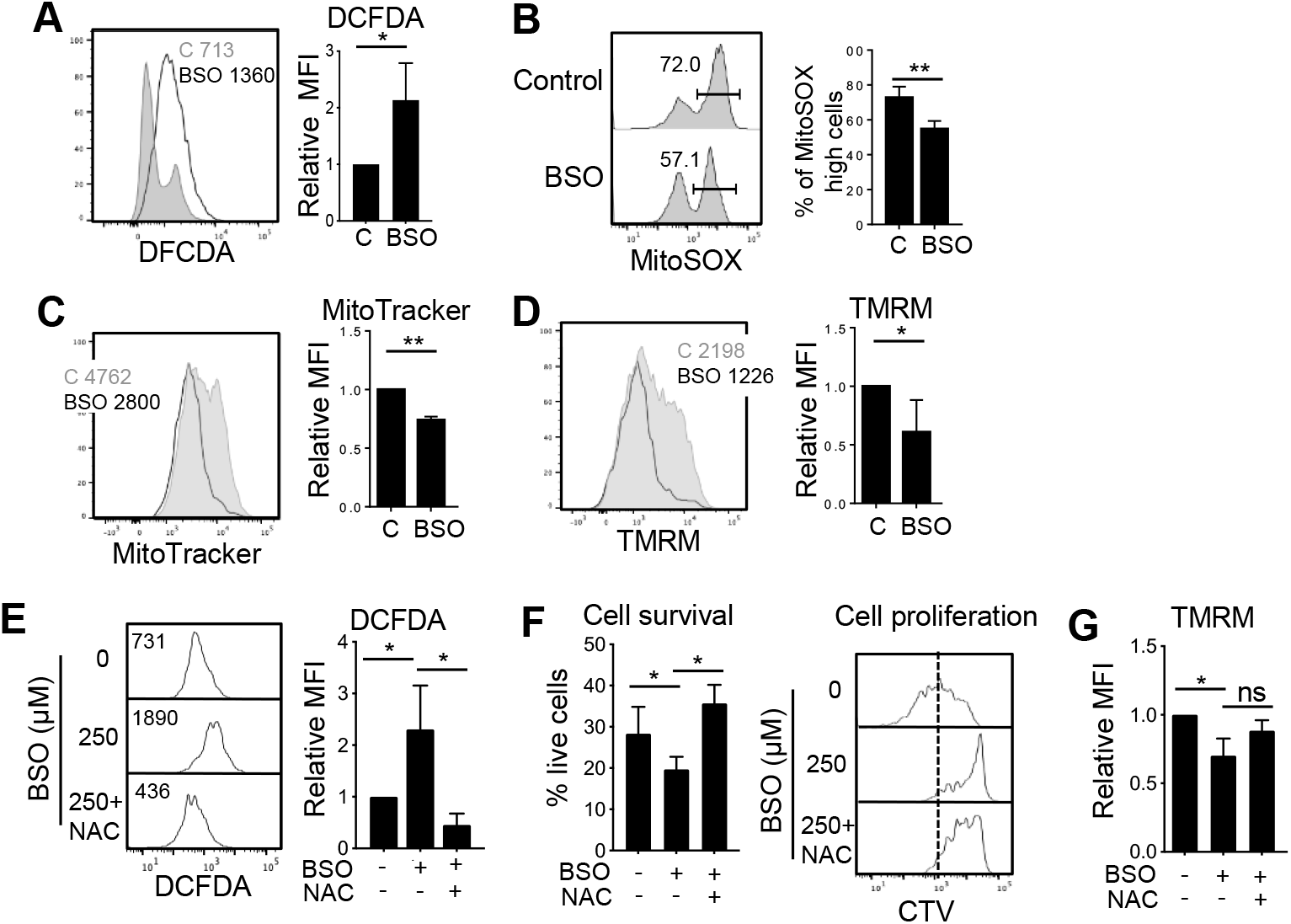
GSH-mediated redox balance is essential for NKT cells homeostasis. Sorted NKT cells from C57BL/6 mice were stimulated for 3 days in the presence or absence of BSO. (A-D) Histograms and graphs show total ROS levels measured using DCFDA (A), mitochondrial ROS measured by MitoSOX (B), mitochondrial mass by MitoTracker (C), and mitochondrial potential by TMRM (D) in activated NKT cells (n=3). (E and F) NKT cells were activated for 3 days in the presence or absence of BSO (250 μM) and NAC (10 mM). Histograms and graphs show total ROS levels (E), cell survival and cell proliferation (F) and mitochondrial potential (G) (n=3). All data are representative of or combined from at least three different experiments. Data are shown as mean ± SEM. *p <0.05, **p<0.01.

These observations suggest that the high levels of cell death in NKT cells after inhibition of GSH synthesis could be due to increased ROS. To test this, we treated cells with the ROS scavenger N-acetyl-cysteine (NAC) to reduce ROS in GSH inhibited cells. We found that NAC restored ROS levels in GSH inhibited cells back to the control levels (Fig. 3E). Interestingly, NKT cell survival was rescued by NAC treatment, whereas cell proliferation (Fig. 3F). The poor proliferation was correlated with the incomplete restoration of mitochondrial membrane potential by NAC (Fig. 3G). Together, these data suggest that NKT cell survival is supported by GSH-mediated redox balance whereas cell proliferation might be supported by GSH-mediated control of mitochondrial function.

### Distinct glutamine oxidation pathways regulate NKT cell effector functions

We have previously shown that glucose availability is critical for NKT cell cytokine production (32). To investigate whether glutaminolysis has any role in cytokine production, we activated NKT cells with and without the GLS inhibitor. Additionally, we activated WT and GLS1 KO NKT cells. GLS activity is critical for IL-4 production in NKT cells, as IL-4^+^ cells were significantly reduced upon CB839 treatment (Fig. S3A). Similarly, both intracellular and secreted levels of IL-4 were lower in GLS1 KO cells than WT NKT cells (Fig. 4A and S3B).

**Fig. 4.**
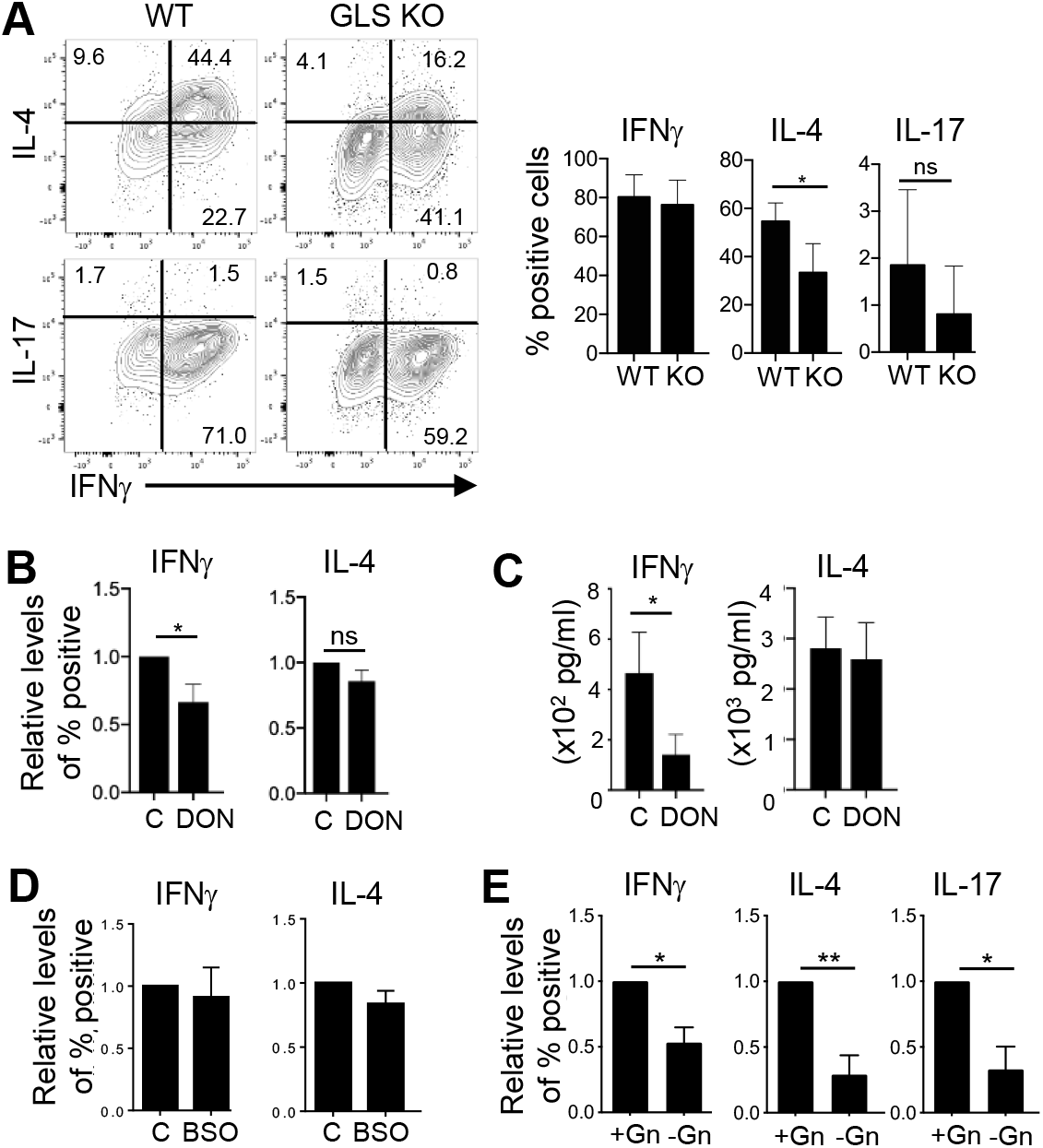
IFNγ and IL-4 production in NKT cells rely on distinct branches of glutamine metabolism. Sorted NKT cells from C57BL/6 mice were stimulated for 3 days in the indicated culture conditions. (A) Representative dot plots show cytokine expression in NKT cells from WT and GLS1 KO mice. Graphs show cumulative data from 3 independent experiments. (B and C) NKT cells were activated in the presence or absence of DON (10 μM). Intracellular cytokine expression (B) and the levels of cytokine secreted into the media by ELISA (C) were measured (n=3). (D) The graph shows cytokine expression in NKT cells stimulated in the presence or absence of BSO (n=3). (E) NKT cells were stimulated with or without glutamine (Gn) and cytokine expression was compared at day 3. Data are shown as mean ± SEM. All data are representative of or combined from at least three independent experiments. *p<0.05, **p<0.01, ns: not significant.

We next asked whether the HBP regulates cytokine production in NKT cells. In contrast to GLS inhibition, inhibition of GFAT1 via DON treatment decreased IFNγ but not IL-4 production (Fig. 4B and 4C). However, inhibition of OGT significantly reduced IFNγ expression but only moderately affected IL-4 expression (Fig. S3C). Overall, our data suggest that *de novo* HBP activity is critical for cytokine production by NKT cells.

ROS seems to be important for NKT cell effector functions at a steady state. However, ROS is decreased upon NKT cell activation (45). To examine whether GSH-mediated redox balance modulates NKT cell effector function, we activated NKT cells in the presence or absence of BSO and measured cytokine expression. Interestingly, cytokine production was not affected by GSH inhibition (Fig. 4D and S3D), even though GSH inhibited cells have higher total ROS levels (Fig. 3A). Similar to GLS inhibition, GDH inhibition led to a dramatic reduction in IL-4^+^ NKT cells but only a slight reduction in IFNγ^+^ NKT cells (Fig. S3E).

Since glutamine fuels both glutaminolysis and the HBP (Fig. 2A), we were interested in investigating the role of glutamine itself in cytokine expression. We stimulated NKT cells in the presence or absence of glutamine and compared the cytokine expression. Glutamine deprivation reduced the expression of IFNγ, IL-4, and IL-17 by NKT cells (Fig. 4E) suggesting a distinct role of glutamine metabolic pathways for cytokine expression in NKT cells.

### Mitochondrial anaplerosis fueled by glutamine-derived αKG is necessary for NKT cell homeostasis and effector function

Glucose can be metabolized through glycolysis to fuel the TCA and produce lactate. Previously, we showed that the expression of PPP genes was significantly higher in NKT cells compared to CD4 T cells (32). Consequently, the levels of glycolytic metabolites were lower in NKT cells than CD4 T cells (Fig. 1A). Additionally, glucose deprivation did not affect NKT cell survival or proliferation, raising the possibility that NKT cells rely primarily on glutamine (32). To determine if NKT cells are addicted to glutamine, we first measured the expression of hexokinase 2 (HK2), which converts glucose into glucose 6-phosphate during the first step of glycolysis. Consistent with our previous finding that CD4 T cells take up more glucose than NKT cells upon activation (32), HK2 expression was also higher in activated CD4 T cells (Fig. 5A). Next, we measured PPP metabolites in NKT and CD4 T cells by LC-MS/MS. Compared to CD4 T cells, NKT cells have notably higher levels of PPP metabolites before activation (Fig. 5B). Additionally, PPP metabolite levels were further increased upon activation (Fig. 5C), suggesting NKT cells are primarily metabolizing glucose via the PPP. Because glutamate-derived αKG plays a key role in mitochondrial anaplerosis for NKT cell survival and proliferation, we investigated whether glucose could fuel mitochondrial activity in the absence of glutamine. To do this, we measured glucose uptake in NKT cells grown under glutamine deprivation conditions. The results showed that NKT cells were not able to efficiently take up glucose under glutamine deprivation conditions (Fig. S4A) and that they preferentially used glutamine to produce ATP (Fig. S4B).

**Fig. 5.**
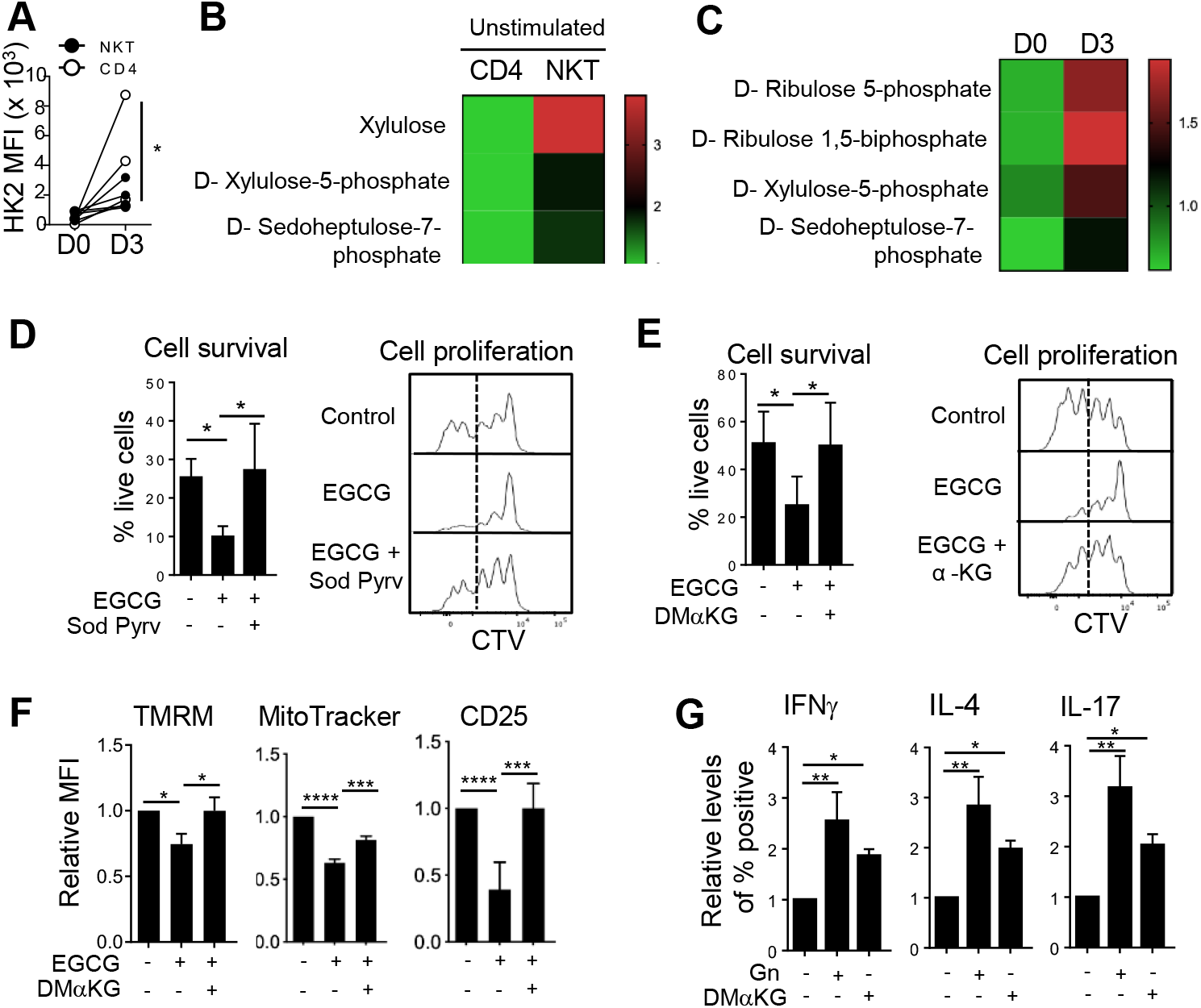
NKT cells exhibit a glutamine-addicted phenotype. (A) The graph shows hexokinase 2 (HK2) expression in NKT cells with and without stimulation (n=3). (B) Heat map shows relative levels of the indicated PPP metabolites in NKT cells compared to CD4 T cells analyzed after LC-MS/MS analysis (n=3). (C) Heatmap shows HBP metabolites in NKT cells with and without stimulation as analyzed by LC-MS/MS (n=3). (D and E) Graphs show cell survival and proliferation of NKT cells stimulated in the presence of EGCG (20 μM) together with sodium pyruvate (1 mM) (D, or with dimethyl alpha-ketoglutarate (DMαKG) (1.5 mM) (E). (n=3). (F) Cells in (E) were used to measure mitochondrial potential, mitochondrial mass, and CD25 expression (n=3). (G) NKT cells were stimulated for 3 days in the presence or absence of glutamine (2 mM) in combination with DMαKG (1.5 mM). Graphs show relative percentages of cytokine positive NKT cells (n=3). All relative levels were calculated using the MFI values of the control as 1. All data are representative of or combined from at least three different experiments. Data are shown as mean ± SEM. *p<0.05. **p<0.01, ***p<0.001, ****p<0.0001. ns: not significant.

Elevated PPP gene expression suggests that glucose is metabolized mainly via the PPP in NKT cells (32) and glucose-derived pyruvate would not be sufficient to supplement mitochondrial anaplerosis in NKT cells. We tested this hypothesis by adding sodium pyruvate during GDH inhibition to see whether the reduced cell survival and proliferation observed after GDH inhibition can be rescued by pyruvate directly. Strikingly, we observed that NKT cell survival and proliferation were restored to control levels after feeding sodium pyruvate to GDH inhibited cells (Fig. 5D).

To further support that glutamate-derived αKG is essential for mitochondrial anaplerosis in NKT cells, we provided dimethyl α-ketoglutarate (DMαKG), a cell-permeable αKG analog, to the culture media in the presence of EGCG. Both cell survival and proliferation were restored to normal levels by αKG supplementation in GDH inhibited NKT cells (Fig. 5E). Additionally, DMαKG partially rescued cell survival under glutamine deprivation conditions (Fig. S4C) as well as cell proliferation in CB839 treated NKT cells (Fig. S4D), suggesting that glutamine-derived αKG is essential to maintain NKT cell survival and proliferation. As expected, DMαKG supplementation not only rescued mitochondrial function and NKT cell activation (Fig. 5F) but also restored cytokine production under either glutamine-deficient culture conditions (Fig. 5G and S4E) or GLS inhibition (Fig. S4F). Similarly, DMαKG corrected the cytokine profiles of GDH inhibited cells (Fig. S4G).

These data demonstrate that NKT cells exhibit lower levels of glycolysis, which is insufficient to provide enough glucose-derived metabolites to the TCA cycle. As a result, NKT cells primarily rely on glutamine to fuel mitochondrial function for their survival, proliferation, and cytokine production.

### Glutaminase is crucial for proper NKT cell responses to *Listeria monocytogenes* infection

To investigate the role of glutamine metabolism in NKT cell-mediated immune responses *in vivo*, we used the *Listeria* infection model. We injected *Listeria monocytogenes* expressing ovalbumin intraperitoneally to WT and GLS1 KO mice. Bacterial load and NKT cell-specific functions were analyzed after 2 days of infection. This time point allows us to study NKT cell-mediated effects on bacterial infection, as CD4 and CD8 T cells are not able to mount an immune response in this short time frame. To examine whether NKT cell metabolic responses are changed after *Listeria* infection, we compared GSH and CD98 expression in WT mice. Both CD98 expression (Fig. 6A) and GSH levels (Fig. 6B) were greatly increased in splenic and hepatic NKT cells in infected mice compared to PBS-injected controls. When bacterial loads were compared, GLS1 KO mice had higher bacterial loads than WT mice in both the spleen and liver (Fig. 6C). Higher bacterial burden correlated with impaired activation of NKT cells from GLS1 KO mice (Fig. 6D). We then asked whether the high bacterial load in GLS1 KO mice is due to slower NKT cell expansion or diminished cytokine expression. We observed that cell proliferation was impaired in NKT cells from GLS1 KO mice in response to bacterial challenge (Fig. 6E), supporting the important role for glutamine metabolism in NKT cell responses. However, like our *in vitro* observations, *Listeria* infection did not significantly affect IFNγ production in NKT cells (Fig. 6F). Overall, GLS-mediated glutaminolysis is essential for NKT cells to mediate protective immune responses against *Listeria* infection.

**Fig. 6.**
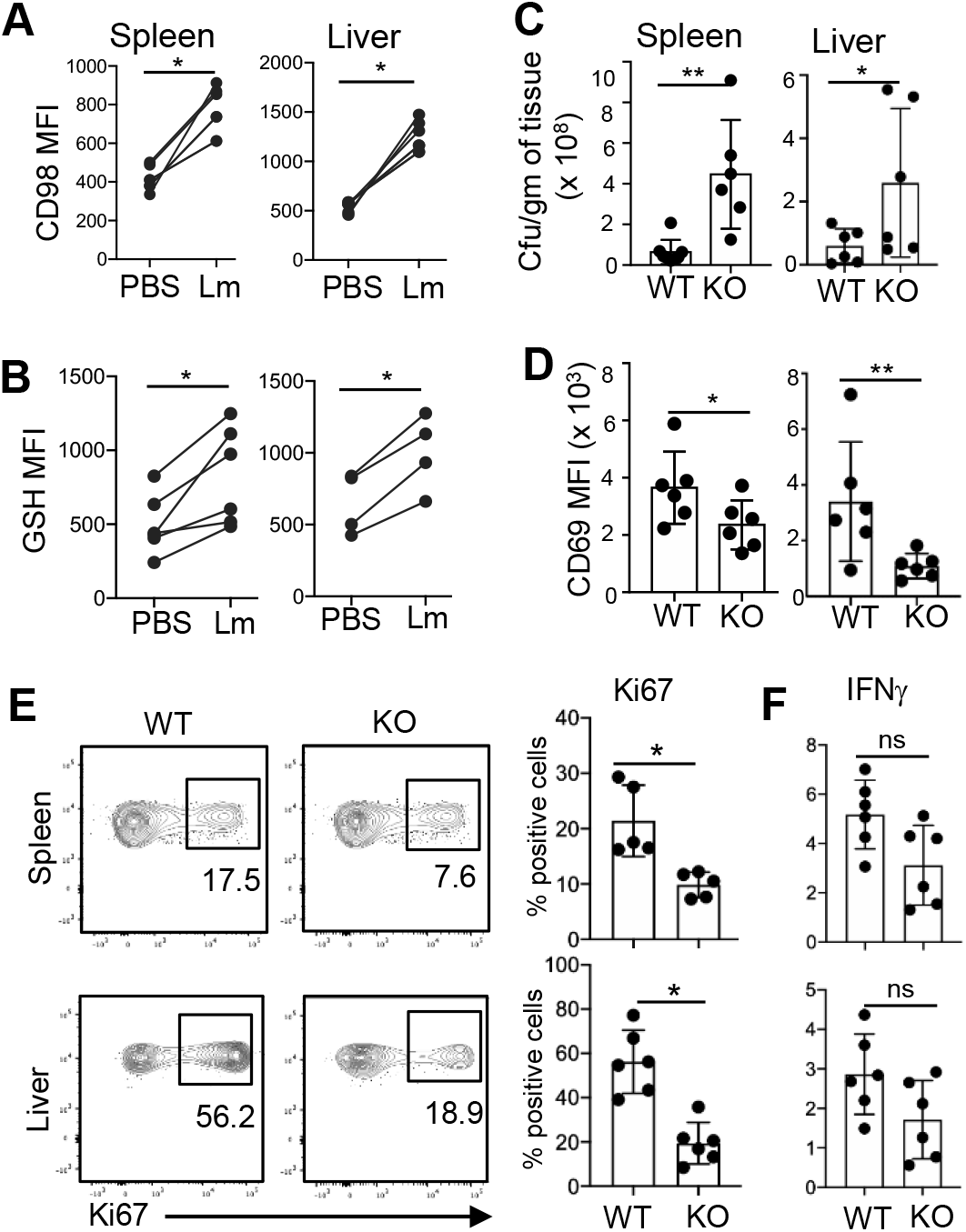
Glutaminolysis is important for NKT cell responses to *Listeria* infection. WT and GLS1 KO mice were injected with either 10^5^ CFU/mouse of LM-Ova (Lm) or PBS intraperitoneally. Two days after infection, spleens and livers were harvested and analyzed for bacterial load, NKT cell proliferation, and IFNγ expression. (A and B) Graphs show levels of CD98 expression and GSH in NKT cells from the spleen (left panel) and the liver (right panel) of PBS- and Lm-injected WT mice (n=6). (C) Graphs show bacterial loads in the spleens and livers of infected WT and GLS1 KO mice (n=6). (D) Graphs show CD69 expression in splenic and hepatic NKT cells from WT and GLS1 KO mice (n=6). (E) Representative dot plots and graphs show cell proliferation as measured by Ki-67 expression in NKT cells from the spleen and liver (n=6). (F) To assess IFNγ expression, total splenocytes were incubated in the presence of Monensin for 2 h to prevent cytokine secretion followed by comparing intracellular expression of IFNγ in splenic (top panel) and hepatic (bottom panel) NKT cells (n=6). Data are shown as mean ± SEM. *p <0.05; **p<0.01. ns: not significant.

### The AMPK-mTORC1 axis regulates NKT cell glutamine metabolism

Studies have linked mTORC1 activation to glutamine addiction in some types of cancer cells (46). Moreover, mTOR signaling is critical for the development and function of NKT cells (25, 47). We have previously shown that mTORC1 activity is enhanced upon NKT cell activation (32). Furthermore, NKT cells stimulated in the presence of high lactate showed reduced mTORC1 activity accompanied by poor proliferation (32). As such, we reasoned that mTORC1 signaling might affect glutamine metabolism in NKT cells. To test this, we used the pharmacological reagent rapamycin to inhibit mTORC1 activity because mTORC1 deficiency compromise NKT cell development (Prevot et al., 2015). We stimulated NKT cells in the presence of rapamycin and examined glycolysis and amino acid transport by comparing the expression of HK2 and CD98, respectively. We also measured glutamine, glutamate, and GSH to study glutaminolysis. mTORC1 inhibition by rapamycin resulted in reduced HK2 expression, CD98 expression, and GSH levels (Fig. 7A-7C). Interestingly, mTORC1 inhibition also reduced proteome O-GlcNAc levels (Fig. 7D), suggesting that mTORC1 is a key regulator of glucose and glutamine metabolism including glycosylation in NKT cells.

**Fig. 7.**
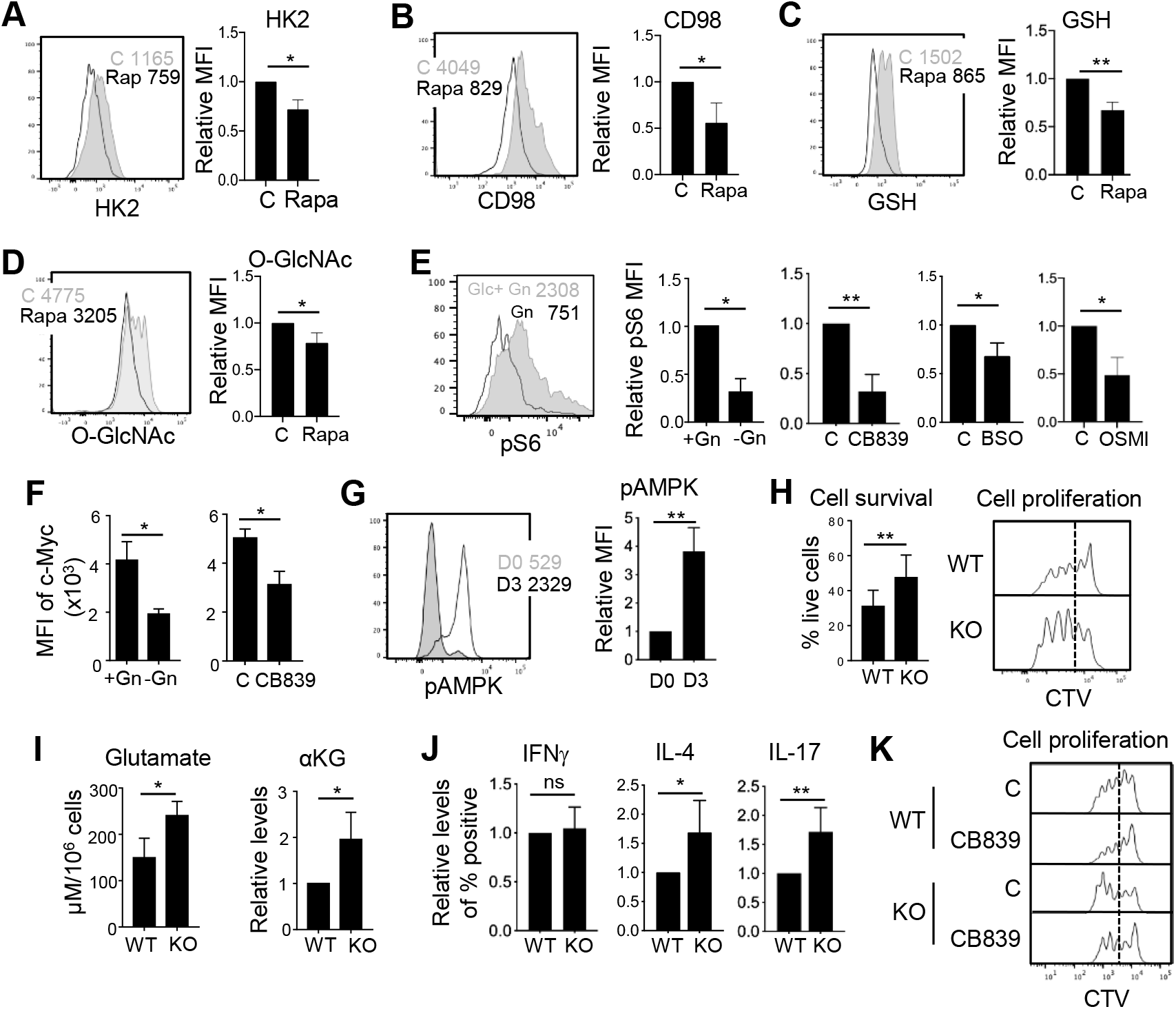
mTORC1-AMPK signaling regulates both glucose and glutamine metabolism in NKT cells. (A-D) Sorted NKT cells were stimulated for three days with and without rapamycin (2 nM). Representative histograms and graphs show relative HK2 expression (A), CD98 expression (B), GSH production (C), and O-GlcNAc levels (D) in stimulated NKT cells (n=3). (E) NKT cells were activated for three days under the indicated conditions and relative expression of pS6^Ser235/236^ was compared (n=3). (F) Graphs show relative expression of c-Myc in NKT cells activated for 3 days in the presence or absence of either glutamine or CB839 (n=3). (G) Relative expressions of pAMPK without (D0) and with (D3) activation are shown (n=3). (H) Cell survival and proliferation of WT and AMPK KO NKT cells after 3 days of activation (n=3). (I) Glutamate and αKG levels were compared between WT and AMPK KO NKT cells after 3 days of activation. (J and K) WT and AMPK KO NKT cells were labeled with CTV and stimulated in the presence or absence of CB839 for 3 days. (J) Graphs show cytokine expression comparison between WT and AMPK KO NKT cells (n=3). (K) Histograms show cell proliferation (n=2). Control levels were set at 1. Data are shown as mean ± SEM. *p<0.05, **p<0.01. ns: not significant.

mTORC1 signaling integrates growth factors and nutrient signals to regulate cell growth. mTORC1 is unresponsive to these signals under amino acid deprivation (48). In particular, glutamine and glutamate are essential for maintaining mTORC1 activity in T cells (7, 49). Therefore, we asked whether glutamine or glutamate availability is necessary for mTORC1 activity in NKT cells. Indeed, both glutamine deprivation and GLS inhibition reduced the phosphorylation of ribosomal protein S6 (pS6), a substrate of mTORC1 (Fig. 7E). Similarly, inhibition of GSH production and OGT activity also decreased mTORC1 activity (Fig. 7E). mTORC1 is known to enhance c-Myc expression (50). We found that c-Myc levels were reduced in NKT cells grown under glutamine deprivation conditions as well as after CB839 treatment (Fig. 7F), indicating that GLS activity regulates mTORC1 signaling in NKT cells. Together, these data suggest that crosstalk between glutamine metabolism and mTORC1 signaling regulates cell proliferation in NKT cells.

The levels of phosphorylated AMPK (pAMPK) increase during primary T cell responses *in vivo* (51), and pAMPK is known to negatively regulate mTORC1 activity in T cells (29). Having observed that mTORC1 promotes glutamine metabolism in NKT cells, we investigated the role of the AMPK. We found that pAMPK levels were greatly increased in stimulated NKT cells compared to unstimulated cells (Fig. 7G). Next, we used T cell-specific AMPK KO mice to test our hypothesis that AMPK deficiency would elevate glutaminolysis, which would have a beneficial effect on NKT cells. AMPK KO mice have no observed defects in conventional T cell (52) or NKT cell development or peripheral maintenance (Fig. S5). We first analyzed activated NKT cell survival and proliferation in WT and AMPK KO mice. AMPK KO NKT cells were more resistant to cell death and proliferated better than WT cells (Fig. 7H). As expected, AMPK KO NKT cells have higher levels of glutamate and αKG (Fig. 7I) and expressed more IL-4 and IL-17 (Fig. 7J), suggesting that glutaminolysis is enhanced in AMPK KO NKT cells. Furthermore, in contrast to WT, AMPK KO NKT cells proliferated efficiently even in the presence of GLS inhibitor (Fig. 7K). Together, this data suggests that the AMPK-mTORC1 signaling axis controls glutamine metabolism in NKT cells.

## Discussion

Glucose and glutamine are the two primary nutrients utilized by highly proliferative cells, including T cells (53). Unlike glucose, glutamine can provide both carbon and nitrogen for anabolic reactions (54). Indeed, glutamine-derived nitrogen is critical for the synthesis of nitrogenous compounds such as nucleic acids, glycosoamino glycans, and non-essential amino acids (55). Here, we demonstrate that glutamine is metabolized via glutaminolysis and the HBP in activated NKT cells to support their survival, proliferation, and effector functions. We also show that NKT cells are glutamine addicted because their low glycolytic rate cannot spare enough glucose-to support the TCA cycle. Moreover, glutamine metabolism seems to be regulated by AMPK-mTORC1 signaling in NKT cells.

Although we comprehensively investigated glutamine metabolism in NKT cells using pathway specific inhibitors, we used T cell specific GLS1 KO mice to confirm GLS inhibition studies. Since these studies were performed on primary NKT cells which are in low abundant than other conventional T cell, the experiments with genetically knocked down of genes using specific siRNAs is quite difficult. In line with this, glutamine tracing experiments were not feasible with primary NKT cells.

Glutamine metabolism is differentially regulated in the various T cell subsets (6, 7). In addition to synthesizing glutamine *de novo*, proliferating cells can acquire glutamine from the extracellular environment to meet their energetic requirements. Resting NKT cells have higher glutamine levels than CD4 T cells, which may explain why they rely on glutamine upon activation for their survival and proliferation. This idea is supported by the fact that GLS1 KO mice exhibit lower NKT cell frequencies in the spleen compared to WT. In addition to glutamine, the levels of other amino acids were also higher in activated NKT cells compared to CD4 T cells. Whether NKT cells have enhanced uptake or increased synthesis of these amino acids from glutamine warrants further investigation. The high rate of glutamine consumption in NKT cells suggests that these cells use glutamine for multiple roles beyond protein synthesis. We found that NKT cells use glutamine in the HBP to modulate protein modification processes like glycosylation. It is important to note that TCA cycle intermediates can regulate epigenetic signatures in activated T cells (6, 7). These metabolites are critical in regulating T helper cell subsets and their cytokine production (56). In corroboration with these facts, inhibiting glutamate oxidation reduced cytokine production by NKT cells, which was rescued by αKG supplementation. Interestingly, NKT cells were observed to rely on glutaminolysis primarily for IL-4 production but depend upon glutamine oxidation from the HBP for IFNγ production. We also showed that NKT cells depend upon GLS to proliferate *in vivo* following *Listeria* infection. However, GLS does not control IFNγ production, indicating that GLS largely controls NKT cell homeostasis. Extending these findings to their *in vivo* relevance, we observed slower NKT cell proliferation and higher bacterial burden after *Listeria* infection in GLS1 KO mice.

Mitochondrial homeostasis is critical for NKT cell development (25, 47). T cell-specific deletion of RISP (T-Uqcr^-^/^-^), a nuclear-encoded protein subunit of mitochondrial complex III, has recently been shown to block NKT cell development (57). Additionally, conventional T cells use glucose to produce lactate and fuel mitochondrial metabolism. These processes are critical for T cell homeostasis and effector function (58). NKT cells have low glucose uptake but high PPP enzyme expression and metabolite abundance. Together, these results suggest that less glucose-derived carbon is oxidized through glycolysis and therefore less is available to support TCA cycling in NKT cells. Based on the reduced availability of glucose-intermediates to fuel respiration, NKT cells instead depend upon glutamine to fuel the TCA cycle via the production of αKG.

ROS can act as signaling messengers as well as positively modify protein structure; however, high concentrations of ROS can lead to cell death (35, 59). Antioxidation via GSH supports activation-induced metabolic reprogramming in T cells (35). Additionally, NKT cells reduce intracellular ROS levels upon activation (45), suggesting that high levels of ROS may be detrimental for activated NKT cells. Our data suggest that glutamine also contributes to GSH synthesis in NKT cells, which is critical for maintaining the redox balance necessary for cell survival. GSH also supports mitochondrial function in NKT cells, and this phenomenon might be due to mTORC1 activation. Further investigation is warranted to shed light on this mechanism.

Lymphocytes must balance a wide range of metabolic pathways to maintain homeostasis after activation. T cells use glutamine-dependent OXPHOS to produce ATP and remain viable in low glucose environments. Because NKT cells have low glycolytic capacity, AMPK is triggered upon activation to regulate glutamine metabolism. AMPK has been reported to regulate mTORC1 in T cells (29). mTORC1 supports glycolysis by directly regulating pathway-specific gene expression in T cells (60). Previously, we have shown that mTORC1 inhibition by rapamycin not only compromised NKT cell survival and proliferation but also reduced glucose uptake (32). Here, we showed that rapamycin treatment negatively affected various steps of glutamine metabolism including glutamine transporter expression, HBP pathway activity, and GSH synthesis.

We report in this manuscript that glutamine fuel the HBP to maintain NKT cell homeostasis and effector function, shedding light on a potential mechanism for NKT survival in the tumor microenvironment (TME). Low availability of glucose in the TME (27, 43) reduces conventional T cell proliferation and cytokine production (61, 62). However, glucose restriction likely does not affect NKT cell homeostasis, as these cells are more dependent upon glutamine. Our findings lead us to propose that NKT cells can be used as an effective immunotherapeutic agent against glucose-reliant tumors.

In conclusion, glutamine oxidation is pivotal for NKT cell survival and proliferation. Because NKT cells display inefficient glycolysis, we predict that they cannot effectively use glucose to fuel mitochondrial metabolism. Glutamine-derived GSH is critical in maintaining redox balance in NKT cells, which is essential for their survival. This study also reveals that NKT cells use different glutamine oxidation pathways for IL-4 and IFNγ production. Moreover, AMPK-mTORC1 signaling regulates glutamine metabolism in NKT cells. Taken together, NKT cells have unique metabolic requirements, and a better understanding of these requirements may contribute to the development of new therapeutic targets to improve T cell-based therapies in the future.

## Materials and Methods

### Mice

Male and female C57BL/6 mice ranging from 8-12 weeks of age were either bred in-house or purchased from The Jackson Laboratory. T cell-specific GLS1 deficient mice (referred to as GLS1 KO) and AMPK deficient mice (referred to as AMPK KO) were generated by crossing GLS1^*fl/fl*^ mice and AMPK^*fl/fl*^ with CD4-Cre expressing mice purchased from The Jackson Laboratory. In all experiments, WT littermates were used as controls. All mice were bred and maintained under specific pathogen free conditions. All animal experiments were performed in accordance with the Institutional Animal Care and Use Committee of the University of Michigan.

### Cell isolation and activation

Primary cell suspensions were prepared from spleens as per standard protocol (45). To sort NKT and CD4 T cells, B cells were excluded from whole splenocytes by incubating with anti-CD19 beads (Miltenyi Biotec) or by using the EasySep™ mouse CD19 positive selection kit (STEMCELL Technologies) as per the manufacturer’s protocol. NKT and CD4 T cells were sorted on the basis of TCR-β and PBS-57 loaded CD1d tetramer expression using a FACS Aria II (BD Biosciences). To study activated NKT cells, cells were stimulated with α-Galactosylceramide (αGalCer; 100 ng/ml) in RPMI 1640 medium supplemented with 10% FBS, 2 mM glutamine, and penicillin/streptomycin at 37°C. For glucose and glutamine deprivation assays, sorted NKT cells were stimulated in glucose- and glutamine-free RPMI 1640 media supplemented with 10% dialyzed FBS (Sigma Aldrich). To inhibit GLS1 activity, CB839 (Sigma Aldrich) was used at 1.5 nM, 3 nM, 250 nM, or 500 nM. To inhibit GDH activity, epigallocatechin-3-gallate (EGCG) (Sigma Aldrich) was used at 10 μM or 20 μM. To inhibit GSH synthesis, L-buthionine-sulfoximine (BSO) (Sigma Aldrich) was used at 250 μM or 500 μM. Rapamycin (Sigma Aldrich) was used to inhibit mTORC1 activity at a concentration of 2nM. 6-diazo-5-oxo-L-norleucine (DON) was used to inhibit O-GlcNAc production at a concentration of 3 μM, 6 μM, or 20 μM. To inhibit OGT1 activity, OSMI-1 (Sigma Aldrich) was used at 10 μM or 20 μM concentrations. For αKG supplementation assays, dimethyl-2-oxoglutarate (DMαKG) (Sigma Aldrich) was used at a 1mM concentration. N-acetyl cysteine (NAC) (Sigma Aldrich) was used at a 1mM concentration as an antioxidant.

### Flow cytometry

The following fluorescently conjugated antibodies were used in the presence of anti-FcγR mAb (2.4G2) for surface and intracellular staining (all from eBioscience): anti-mouse TCRβ (H57-597) Pacific Blue/APC, PBS-57 loaded CD1d tetramer APC/PE/Pacific Blue, anti-mouse CD4 APC-Cy7, anti-mouse IFNγ PE/FITC, anti-mouse IL-4 PE-Cy7, and anti-mouse IL-17 PerCP-Cy5.5. Ki-67 PerCP-Cy5.5 staining was used to measure *in vivo* cell proliferation after *Listeria* infection. Dead cells were excluded by staining with LIVE/DEAD™ Fixable Yellow Dead Cell Stain Kit (405 nm excitation) (Invitrogen).

For intracellular cytokine expression, activated cells were re-stimulated with of PMA (50 ng/mL, Sigma Aldrich) and of Ionomycin (1.5 µM, Sigma Aldrich) in the presence of Monensin (3 µM, Sigma Aldrich) for 4 h. Cells were then stained for surface antigens and intracellular cytokines according to manufacturer’s instructions (BD Biosciences). For intracellular staining of phosphorylated ribosomal protein S6 (pS6^Ser235/236^) (Cell Signaling), HK2 staining (EPR20839) (Abcam), and O-GlcNAc (BD Biosciences), cells were permeabilized using 90% methanol. Cells were then incubated with pS6 or HK2 antibody for 1 h and O-GlcNAc antibody for 20 min at RT in the dark in cytoplasmic permeabilization buffer (BD Biosciences). Nuclear permeabilization buffer was used for c-Myc staining. For cell proliferation, NKT cells were labeled with 5 µM CellTrace™ Violet (CTV) (Invitrogen) in 1X PBS containing 0.1% BSA for 30 min at 37°C. Cells were stimulated as indicated and analyzed by flow cytometry on day three post-stimulation for CTV dilutions. Cells were acquired on a FACS Canto II (BD Biosciences). The data was analyzed using FlowJo (TreeStar software ver. 10.7.1).

### Analysis of metabolic parameters

To measure metabolic parameters, activated NKT cells (1 × 10^5^) were incubated with different reagents as indicated in the fig. legends. To measure mitochondrial parameters, cells were incubated with 60 nM of the potentiometric dye tetramethylrhodamine methyl ester perchlorate (TMRM) (Invitrogen), 30 nM MitoTracker™ Green (Invitrogen), and 2.5 µM MitoSOX (Invitrogen) for 30 min at 37°C in RPMI 1640 complete media. To measure total cellular ROS, 1 × 10^5^ activated NKT cells were incubated with 1 mM 2’,7’-dichlorodihydrofluorescein diacetate (H_2_DCFDA) (Invitrogen) in RPMI complete media for 30 minutes at 37°C. To measure glucose uptake, cells were incubated in 2-(N-(7-nitrobenz-2-oxa-1,3-diaxol-4-yl) amino)-2-deoxyglucose (2-NBDG) (Invitrogen) (20 µM) for 1 h or as indicated at 37°C in glucose-free RPMI 1640 media containing 10% dialyzed FBS. To measure GSH, cells were stained using an intracellular glutathione detection assay kit (Abcam) for 20 min at 37°C in RPMI 1640 complete media. Cells were stained for surface antigens and acquired on a FACS Canto II (BD Biosciences).

### ATP, glutamine/glutamate, αKG, and lactate assays

CellTiter-Glo® Luminescent Cell Viability reagent (Promega) was used for ATP measurement. Intracellular lactate levels were measured using a plate-based fluorometric measurement kit (Cayman Chemicals), while glutamate levels were measured using Glutamine/Glutamate-Glo™ Assay kit (Promega). αKG was measured using a colorimetric assay kit (Sigma Aldrich). All kits were used according to manufacturer’s instructions.

### Metabolite measurements

Cell lysate was prepared from resting NKT and CD4 T cells as well as stimulated NKT cells (5 × 10^5^ cells per replicate) by incubating the cells with 80% methanol and following a series of vigorous mixing steps. Media was mixed with 100% methanol and vigorously vortexed. Cells and media were spun down at maximum speed for 10 min at 4°C to remove membranous debris, and the lysate was collected for drying using a SpeedVac. Following drying, the lysate was reconstituted using 50/50 methanol/water for mass spectrometry-based metabolomics analysis using an Agilent 1290 Infinity II UHPLC combined with an Agilent 6470 QQQ LC/MS.

### RT Gene PCR assay

Total RNA was isolated from unstimulated and stimulated NKT cells using a RNeasy Plus mini kit (Qiagen) according to manufacturer’s instructions. PCR Array was performed according to the manufacturer’s instructions (Qiagen) using Applied Biosystem’s 7900HT Sequence Detection System. Fold changes were calculated from ΔCt values (gene of interest Ct value - an average of all housekeeping gene Ct values) using the ΔΔCt method. Gene expression of target genes was normalized to β-actin.

### *Listeria monocytogenes* infection

*Listeria monocytogenes* expressing ovalbumin (LM-Ova strain 10403s) was grown in BHI broth media. Bacteria in a mid-log phase were collected for infection. GLS1 KO and WT littermate mice were injected intraperitoneally with either 200 μL of sterile 1X PBS alone or 200 μL of 1X PBS containing 10^5^ CFU/mouse of LM-Ova. On day two post-infection, the bacterial burden was enumerated from homogenized spleen and liver samples by culturing serially diluted samples on LB agar plates and performing CFU determination. Intracellular cytokine expression by NKT cells was measured as described above.

### Statistical analysis

All graphs and statistical analyses were prepared using Prism software (Prism version 8; Graphpad Software, San Diego, CA). For comparison among multiple groups, data were analyzed using one-way ANOVA with the multi-comparison post-hoc test. Unpaired and paired Student’s t-tests were used for comparison between two groups. P < 0.05 was considered statistically significant.

## Acknowledgements

We would like to thank Ms. Chauna Black for maintaining our mouse colony and performing genetic screening of all mice. We thank Dr. Mary O’Riordan (University of Michigan) for providing the 10403s LM-Ova strain of *Listeria monocytogenes*. Lastly, we acknowledge the National Institutes of Health Tetramer Facility for providing the CD1d tetramers necessary to study NKT cells.

This work was supported in part by National Institutes of Health Grants R01 AI121156 and R01 AI148289 (to C-H.C.).

C.A.L. was supported by the NCI (R37 CA237421, R01 CA248160). Metabolomics studies performed at the University of Michigan were supported by NIH grant DK 097153, the Charles Woodson Research Fund, and the UM Pediatric Brain Tumor Initiative.

## Author Contributions

A.K. planned and conceived the project, performed experiments, interpreted the data, and wrote the manuscript. C-H.C. secured funding, supervised the project, interpreted the data, and assisted in writing the manuscript. E.L.Y. performed experiments and helped in writing the manuscript. A.A. and L.Z. performed experiments. C.A.L. provided reagents and feedback as well as analyzed the data. All the authors have proofread the manuscript.

