## Supplementary figures for "Glutamine metabolism enables NKT cell homeostasis and function through the AMPK-mTORC1 signaling axis"

This PDF file includes Supplemental Figures 1-5.

**Figure S1**

**A**

| Pathway Name | Match Status | p | -log(p) | Impact |
|---|---|---|---|---|
| Alanine, aspartate and glutamate metabolism | 9/24 | 9.7215E-6 | 11.541 | 0.7289 |
| Purine metabolism | 14/68 | 7.6799E-5 | 9.4743 | 0.304 |
| Pyrimidine metabolism | 10/41 | 2.0564E-4 | 8.4894 | 0.22992 |
| Phenylalanine, tyrosine and tryptophan biosynthesis | 3/4 | 0.0010784 | 6.8323 | 1.0 |
| D-Glutamine and D-glutamate metabolism | 3/5 | 0.0025663 | 5.9653 | 1.0 |
| Arginine and proline metabolism | 8/44 | 0.0067465 | 4.9987 | 0.23187 |
| Taurine and hypotaurine metabolism | 3/8 | 0.012405 | 4.3896 | 0.42857 |
| Phenylalanine metabolism | 3/11 | 0.031594 | 3.4548 | 0.64815 |
| Aminoacyl-tRNA biosynthesis | 9/69 | 0.034417 | 3.3692 | 0.0 |
| Nicotinate and nicotinamide metabolism | 3/13 | 0.049733 | 3.0011 | 0.20833 |
| Butanoate metabolism | 4/22 | 0.052807 | 2.9411 | 0.02899 |
| Pantothenate and CoA biosynthesis | 3/15 | 0.0719 | 2.6325 | 0.02041 |
| Glycolysis or Gluconeogenesis | 4/26 | 0.088274 | 2.4273 | 0.13904 |
| Amino sugar and nucleotide sugar metabolism | 5/37 | 0.09311 | 2.374 | 0.23554 |
| Cysteine and methionine metabolism | 4/27 | 0.098579 | 2.3169 | 0.22557 |
| Nitrogen metabolism | 2/9 | 0.11559 | 2.1577 | 0.0 |
| Vitamin B6 metabolism | 2/9 | 0.11559 | 2.1577 | 0.07843 |
| Citrate cycle (TCA cycle) | 3/20 | 0.14224 | 1.9502 | 0.09365 |
| Fructose and mannose metabolism | 3/21 | 0.15837 | 1.8428 | 0.12754 |
| Valine, leucine and isoleucine biosynthesis | 2/11 | 0.16221 | 1.8189 | 0.0 |

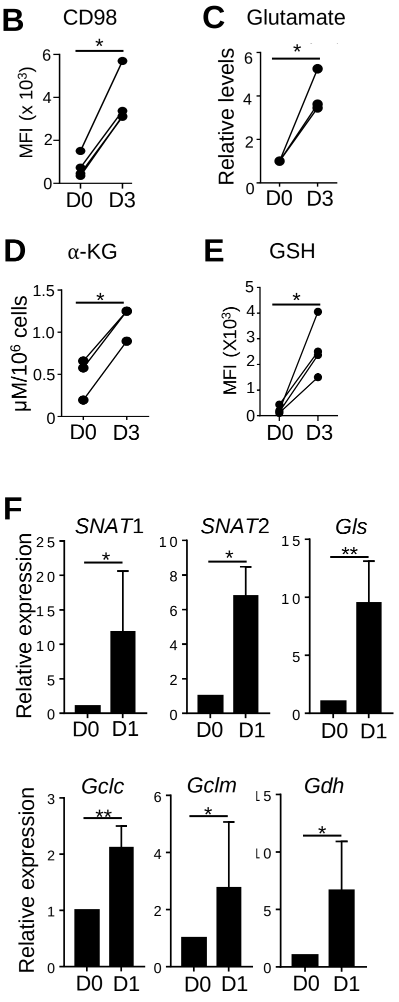


**Figure S1 (Related to Figure 1).** (A) Freshly sorted NKT and CD4 T cells from C57BL/6 mice were subjected to metabolomic analysis through LC-MS/MS. Table shows a summary of the most upregulated metabolic pathways in NKT cells compared to CD4 T cells (n=3). (B-E) Graphs show the levels of CD98 expression (B), glutamate (C), α-KG (D), and GSH (E) in NKT cells with and without stimulation (n=3). D0 sample values were set at 1 to calculate the relative levels of glutamate. All data are representative of or combined from at least three replicates. (F) Graphs show expression of the indicated genes in NKT cells stimulated without (D0) or with (D1) αGalCer (100 ng/mL) (n=3). Data are shown as mean ± SEM. *$p<0.05$, **$p<0.001$.

**Figure S2**

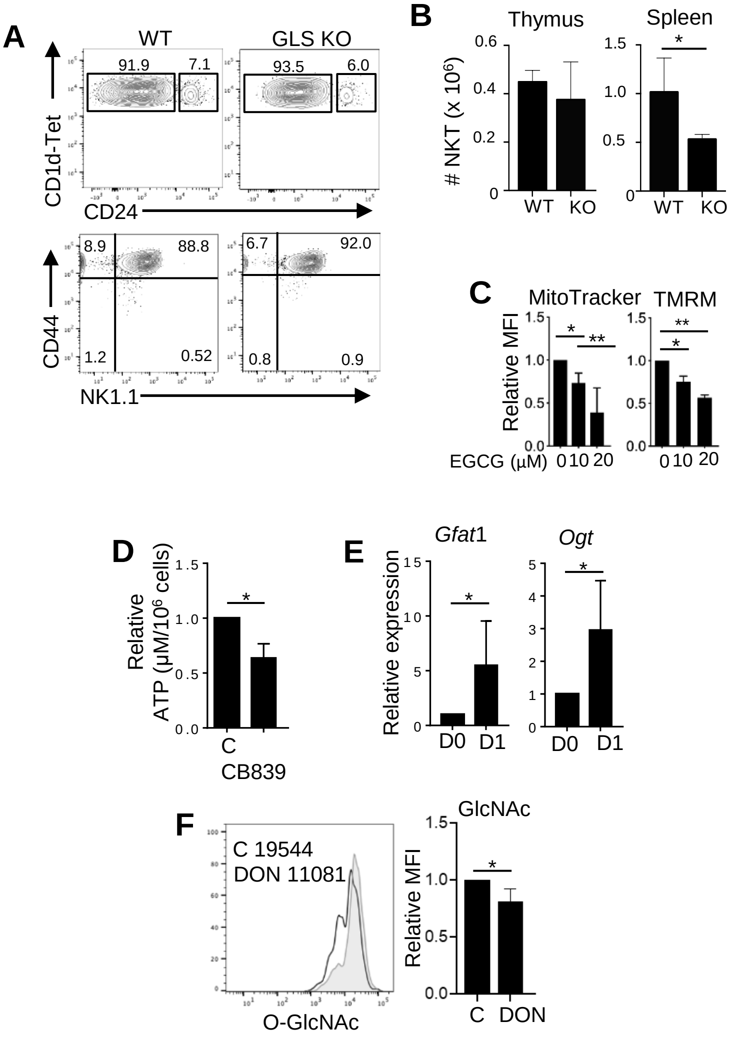


**Figure S2 (Related to Figure 2).** (A) Thymic NKT cells from WT and GLS KO mice were analyzed for expression of CD24 to identify Stage 0 ($CD24^+$) cells as well as CD44 and NK1.1 to identify stage 1 ($CD44^-NK1.1^-$), stage 2 ($CD44^+NK1.1^-$), and stage 3 ($CD44^+NK1.1^+$) cells (n = 3). (B) Graphs show NKT cell numbers in thymi (left panel) and spleens (right panel) of WT and GLS KO mice (n=5). (C) Graphs show mitochondrial mass and potential using MitoTracker and TMRM staining, respectively (n=3). (D) Graph shows relative ATP production in NKT cells activated with and without CB839 (250 nM) for three days. (E) Graphs show relative expression of the indicated genes in resting and stimulated NKT cells (n=4). F) Sorted NKT cells were activated with and without DON (6 μM). On day3, NKT cells were stained for GlcNAc level (n=3). Data are shown as mean ± SEM. *p<0.05, **p<0.01.

#### Figure S3

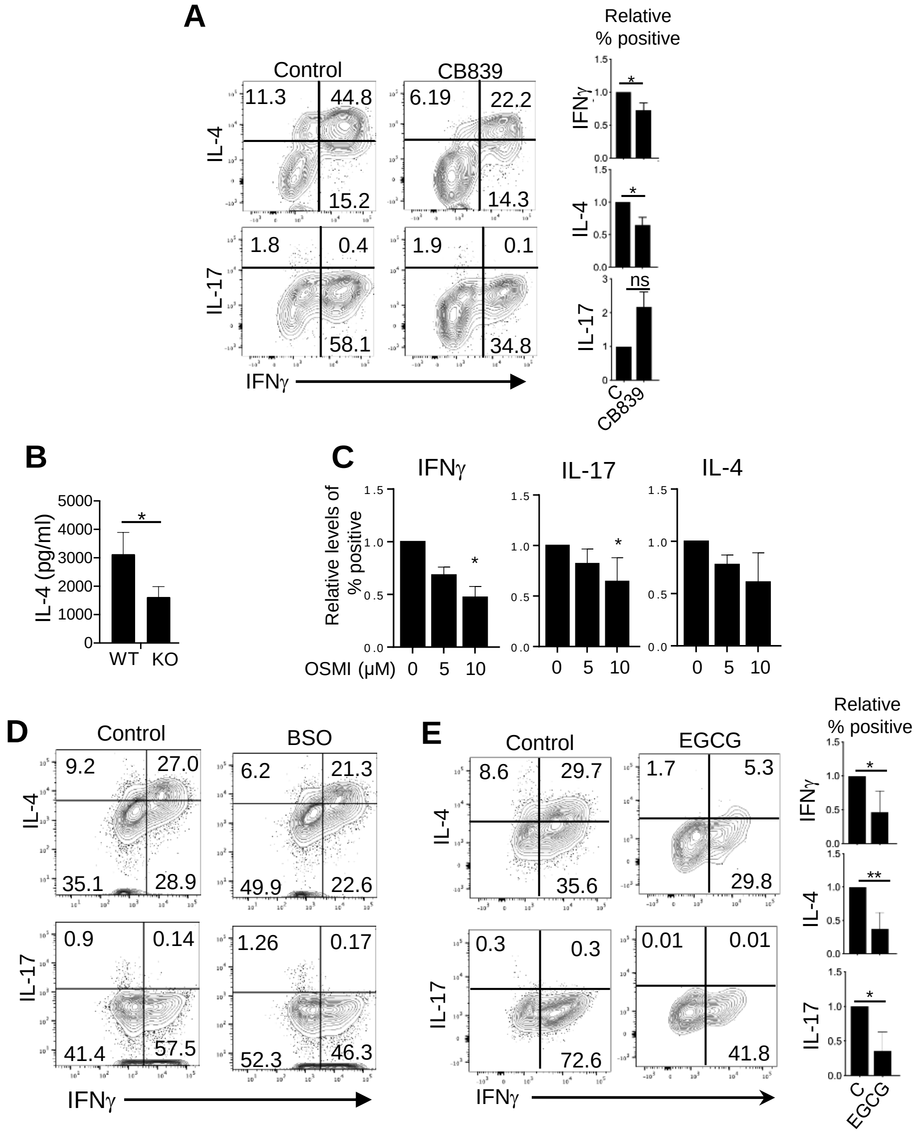


**Figure S3 (Related to Figure 4).** Sorted NKT cells from C57BL/6 mice were stimulated as described in Figure 1E. At day 3 post-activation, cells were re-stimulated with PMA/Ionomycin and Monensin for 4 h followed by intracellular cytokine staining. (A) Cytokine expression in NKT cells activated with and without CB839 (250 nM) (n=3). (B) Graph shows the level of IL-4 secreted into the media by WT and GLS KO NKT cells as measured by ELISA (n=3). (C) Cytokine expression in NKT cells activated with (5 μM and 10 μM) and without OSMI (n=3). (D and E) Cytokine expression in NKT cells activated with or without BSO (250 μM) (D) or EGCG (20 μM) (E) (n=3). Data are shown as mean ± SEM. *$p<0.05$, **$p<0.01$.

### Figure S4

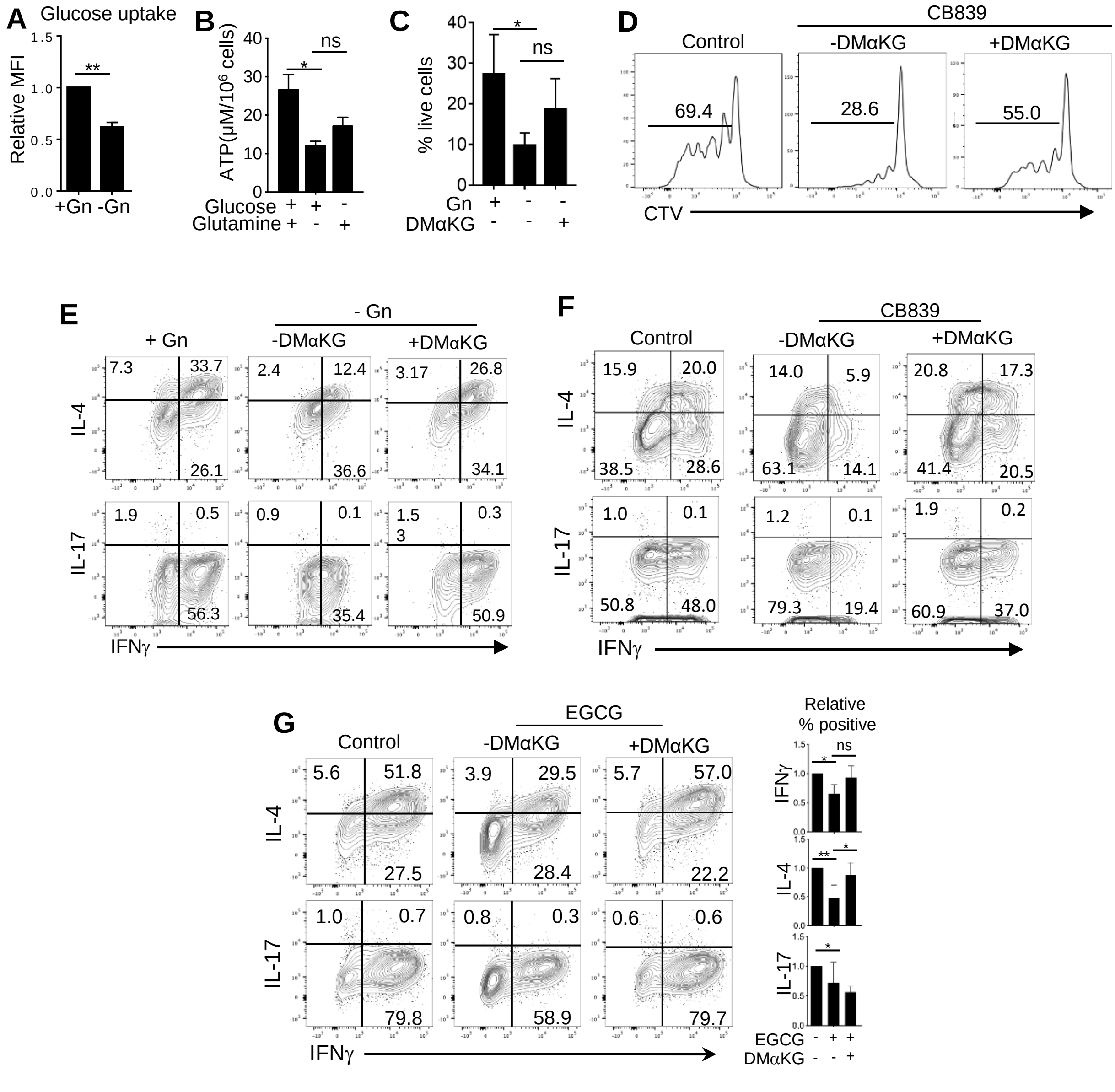


**Figure S4 (Related to Figure 5).** Sorted NKT cells from C57BL/6 mice were stimulated as described in Figure 1E under the indicated culture conditions. (A) Graph shows relative glucose uptake in NKT cells as measured by the mean fluorescent intensity (MFI) of 2-NBDG (n=3). (B) Graph shows ATP production in NKT cells activated under the specified culture conditions (n=3). (C-G) Sorted NKT cells were stimulated with and without glutamine (Gn) (2 mM), CB839 (250 mM), or EGCG (20 μM) in the presence or absence of DMαKG (1.5 mM). (C) Graph shows NKT cell survival (n=3). (D) Histograms show cell proliferation as measured by CTV in NKT cells (n=3). (E-G) Representative dot plots show the cytokine expression in NKT cells treated as noted (F, n=2) (E and G, n=3). Data are shown as mean ± SEM. *p<0.05, **p<0.01.

**Figure S5**

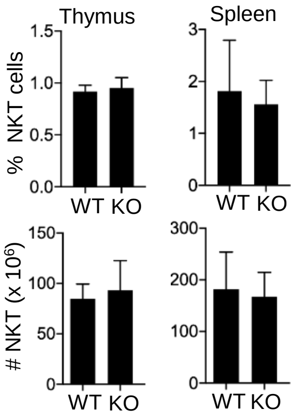


**Figure S5 (Related to Figure 7).** Total cells from the thymi and spleens of WT and AMPK KO mice were counted. Cells were stained for surface markers and analyzed by flow cytometry for NKT cell frequencies. Graphs show percentages (upper panels) and numbers (bottom panels) of NKT cells in the thymi and spleens of WT and AMPK KO mice (n = 3).
